# The Microbe Directory v2.0: An Expanded Database of Ecological and Phenotypical Features of Microbes

**DOI:** 10.1101/2019.12.20.860569

**Authors:** Maria A. Sierra, Chandrima Bhattacharya, Krista Ryon, Sophie Meierovich, Heba Shaaban, David Westfall, Rawhi Mohammad, Katerina Kuchin, Ebrahim Afshinnekoo, David C. Danko, Christopher E. Mason

## Abstract

The Microbe Directory (TMD) is a comprehensive database of annotations for microbial species collating features such as gram-stain, capsid-symmetry, resistance to antibiotics and more. This work presents a significant improvement to the original Microbe Directory (2018). This update adds 68,852 taxa, many new annotation features, an interface for the statistical analysis of microbiomes based on TMD features, and presents a portal for the broad community to add or correct entries. This update also adds curated lists of gene annotations which are useful for characterizing microbial genomes. Much of the new data in TMD is sourced from a set of databases and independent studies collating these data into a single quality controlled and curated source. This will allow researchers and clinicians to have easier access to microbial data and provide for the possibility of serendipitous discovery of otherwise unexpected trends.

## Introduction

The revolutionary advances in sequencing technologies have ushered a large quantity of microbiomes and metagenomics that continue to expand our archives of microbial sequences and genomes (23). Consequently, databases are critical to record, store, maintain, and share information of microbes (28). However, most of the available microbial databases are focused on a particular type of organism pertaining to Bacteria, Fungi or Virus, specially those affecting human health (5; 25; 26), or specific microbial characteristics such as the presence of specific genes or proteins (9; 10; 13; 18; 27). Additionally, most of these databases are not designed to retrieve organisms from different domains sharing the same metadata. For example, when identifying an extremophile that thrives at high temperature, there are archaeal, bacterial, and fungal species that can match that description. There currently was not a database that not only had information across all these domains, but offered the user the ability to filter through either by metadata, taxonomy, and microbiomes.

Thus, The Microbe Directory (TMD) emerged in 2018 as a necessity of a free, user-friendly, and practical database that could be used for a variety of users, from students to scientists and clinicians (22). The first version required 46 trained volunteers to curate metadata for 7,500 microbial species describing eleven microbial characteristics with a binary answer. This second version compiles 16 databases and independent studies into one single inventory, recovering more than 60,000 microbial species. The inventory now includes an increased number of microbial parameters and a classification depending on the microorganism’s domain, due to large genotypic, phenotypic, and ecological differences. Answers to parameters are no longer reduced to a binary format, instead, we have expanded the answer choices in order to be more descriptive. Finally, users are able to easily download or upload information through the web interface, allowing continuous enrichment of the database. The Microbe Directory keeps growing to become a world-free reliable database. TMD version 2 has evolved from a manually curated inventory to a code-curated database leading to a greater number of microbial species, metadata and analysis.

## The Database

The Microbe Directory has expanded in order to address the demand for an all encompassing microbial database, a natural progression that has led to TMD version 2.0. The original version of TMD kept all taxa in a single table. This posed certain challenges and the risk of possible misleading annotations, as certain features were not applicable to all microorganisms. For instance, describing capsid symmetry pertain to viruses while gram stain is specific to bacteria. However, certain features are applicable to both bacteria and fungal species, such as whether they are spore forming or produce biofilms. Thus, to address this issue, the expanded version of TMD divides microbes into one of three tables based on their taxonomic domain: Bacteria and Archaea, Viruses, and Eukaryotes (Figure 1). Each of the three tables contains unique features relevant to that domain reducing possibly misleading annotations. We further describe the processes for collating and cleaning the constituent databases used to build TMD v2.0.

**Figure 1:**
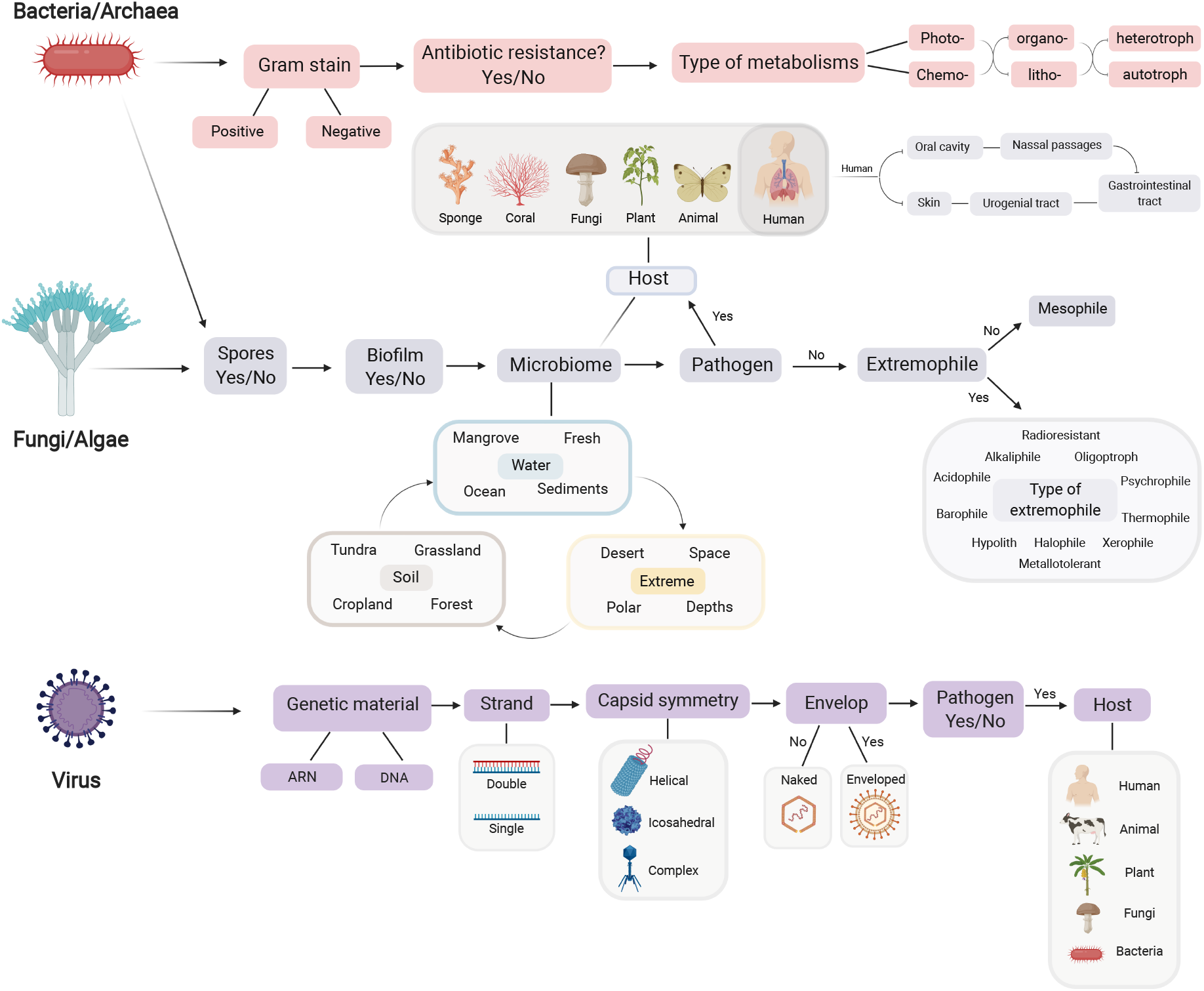
Domain based schema for The Microbe Directory v2.0. The schema above depicts the three domains that taxa in TMD v2.0 are classified into, as well as the major features found in each domain. Certain features are interconnected across multiple domains (e.g. spore forming, biofilms, pathogenicity)

### Microbial features

TMD contains a number of phenotypic characteristics for microbes (Table 1). For example, TMD classifies microbes as mesophiles or extremophiles, and defines mesophiles as microorganisms that grow at an optimal pH and between 14 and 21°C (57 and 70°F). Extremophiles are classified by several types depending on their environmental niche, and these types are not necessary exclusive. *Thermococcus barophilus*, which lives inside hot rocks deep under Earth’s surface, is classified as an extremophile which is both thermophilic and barophilic. The full list of all features with extended descriptions of relevant criteria may be found in the associated web resources.

**Table 1:** Database features and descriptions in the Microbe Directory

| Feature | Description |
| --- | --- |
| Animal Pathogen | Is the microbe a pathogen for any animals? |
| Antimicrobial Susceptibility | Is the microbe susceptible to any antimicrobials? |
| Biofilm Forming | Does the microbe form biofilms? |
| Disease | Is the microbe tied to a specific disease? |
| EMP Site | Sites derived from the Earth Microbiome Project. |
| Extreme Environment | Is the microbe found in extreme environments? |
| Gram Stain | What is the Gram Stain? |
| Host Name | What species host this microbe? |
| Human Commensal | Is the species a known Human commensal? |
| KEGG Disease | KEGG disease ontology |
| KEGG Genome | KEGG genome ontology |
| MetaSUB Site | Sites and materials derived from MetaSUB. |
| Optimal Ph | Optimal pH for the microbe to grow. |
| Optimal Temperature | Optimal temperature for the microbe to grow? |
| Pathogenicity | The COGEM pathogenicity rating of the microbe. |
| Plant Pathogen | Is the microbe known to be pathogenic to plants |
| Spore Forming | Can the microbe form spores? |
| Halotolerance | Is the microbiome halophilic? |
| Soil | Types of soils where this microbe has been found. |
| Tara Ocean | Oceanic sites where this microbe was found. |
| High pH | Can the microbe survive in high pH? |
| HMP Body Site | Sites derived from the Human Microbiome Project. |
| Low pH | Can the microbe survive in low pH? |
| Specific AMR | Specific antimicrobial vulnerabilities. |
| Virus Lineage | Taxonomy of the virus including type of virus. |

### Annotation of features

Taxonomy and microbe names used in TMD are derived directly from the NCBI Taxonomy. All microbes are identified by their current (at time of publication) listed scientific name and numeric taxonomic ID. Unlike the original version of The Microbe Directory all taxonomic ranks may be included in this version of TMD. To annotate features at different ranks we employ a system of infilling based on the taxonomic tree. Thus, if a high taxonomic rank (i.e. Phylum) has a particular trait (e.g. Gram-Stain) all taxa within that rank are assumed to share that trait unless explicitly noted otherwise. Conversely, if all the taxa of a particular rank share a single value for a trait this value is given to the higher rank as well. For ecological locations, particularly those derived from amplicon-based (16S, 18S, or ITS) studies, it is not always possible to derive low level taxa (i.e. Species) from the data. In this case if a particular species belongs to a group (i.e. Genus) that was observed but could not be resolved, we note that the lower rank (Species) belongs to a larger group (Genus) which was observed, but that the lower rank itself was not directly observed. In general we limit this propagation to genus and lower taxonomic ranks.

### Microbiome prevalence

In addition to microbial phenotypic characteristics, TMD v2.0 also describes ecological characteristics of microbes, principally biomes where those microbes are found. This data is collated by probing published taxonomic lists from several large survey studies including Meta-SUB (7) and the Earth Microbiome Project (EMP) (24) (Table 2). We calculated the prevalence of each microbe in each biome (fraction of samples where the microbe was found) and assigned it to one of 5 bins: 0% never observed, up to 25% rarely observed, up to 70% fairly observed, up to 99% mostly observed, and always observed, (Figure 2).

**Table 2:** Databases and studies used for The Microbe Directory v.2.0

| Name | Description | Organism of study | Datasource |
| --- | --- | --- | --- |
| GBIF | Global Biodiversity Information Facility.(GBI) | Bacteria, Virus, Fungi | Database |
| Virus-Host DB | Relationships between viruses and their hosts. (15) | Virus | Database |
| FungiDB | Genome sequence and annotations (2) | Fungi | Database |
| HaloDom | Halophilic organisms (14) | Bacteria, Archaea, Eukarya | Database |
| MyCoPortal | Mycology collections data Portal (16) | Fungi | Database |
| ISHAM Barcoding | ITS reference DNA barcoding fungal database (12) | Fungi | Database |
| RefSoil | Genomes of organisms from the soil (6) | Bacteria and Archaea | Database |
| TARA Ocean | Global Ocean Microbiome (3) | Bacteria, Archaea, Eukarya, Virus | Study |
| MetaSUB | Metagenomics of Subways and Urban Biomes (7) | Bacteria, Archaea, Virus, Eukarya | Study |
| EMP | Earth Microbiome Project (24) | Bacteria, Archaea, Eukarya | Study |
| Rain Forest | Microbiome of lowland tropical rain forest (17) | Bacteria | Study |
| Phyllosphere | The ecology of bacterial communities on tree leaves (19) | Bacteria | Study |
| Airbone | Microbial Communities at High-Elevations (4) | Bacteria | Study |
| Soil pH | Soils microbiome across pH and time (20) | Bacteria and Fungi | Study |

**Figure 2:**
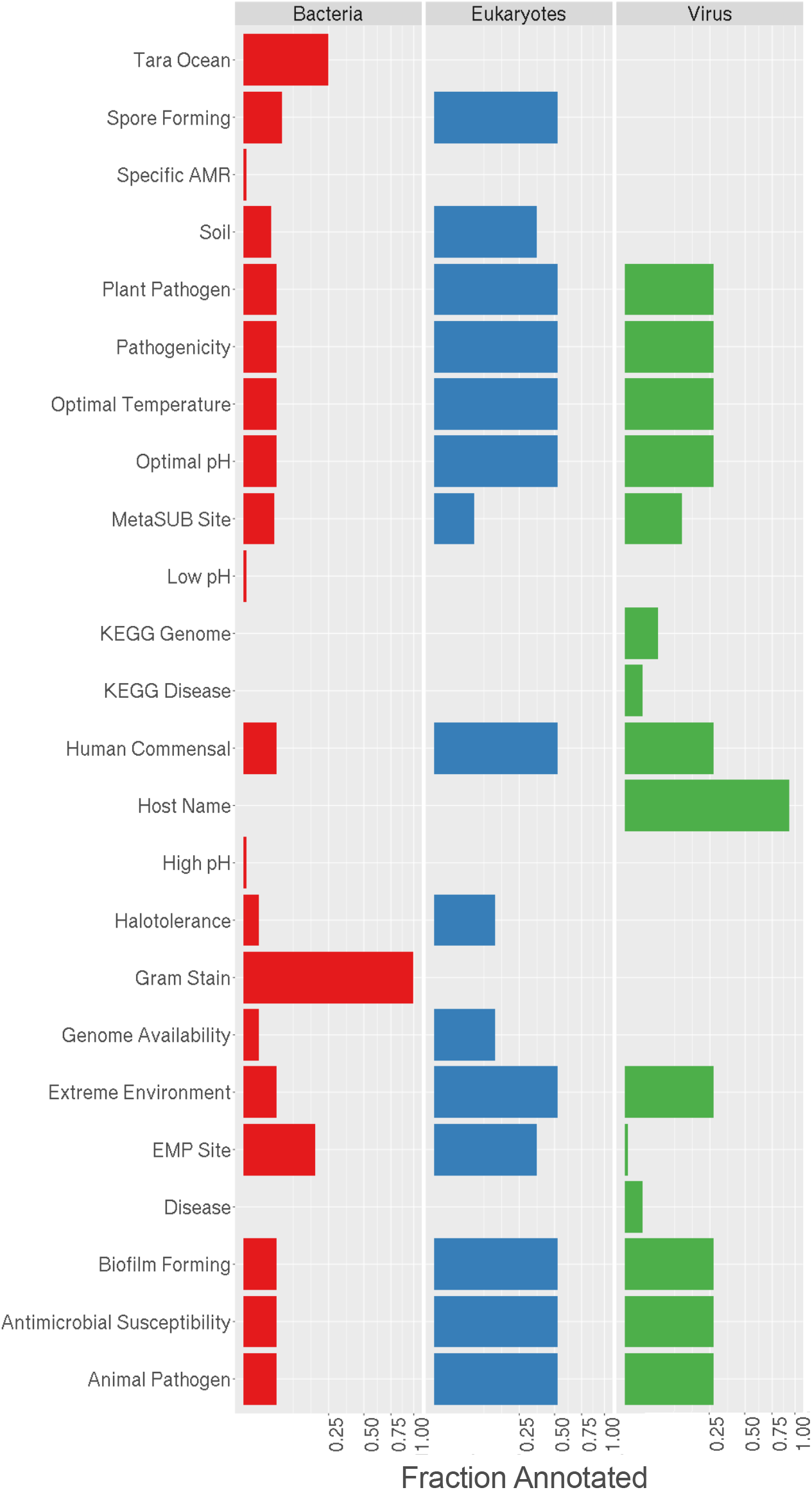
Fraction of features in the Microbe Directory v2.0. Microbial features are filled for the domains. Not all features apply to each domain. Each row represents a particular feature in the Microbe Directory. Not all annotations are present for all taxa and the length of each colored bar represents the fraction of annotated taxa.

### Annotated Gene Sets

In addition to information on microbes, TMD v2.0 contains curated lists of microbial genes with particular functions. The gene descriptions are derived from PROKKA (21), annotations of genomes and are grouped into lists of terms sharing similar high level function. The current gene lists include: Biocide Resistance, Drug Resistance, Mobility, DNA Repair, and Sporulation. These lists were produced by manual curation. The addition of this information provides further utility of TMD. For instance, TMD v1.0 offered researchers to define the characteristics and features of the organisms they characterized in their microbiome samples (i.e. what percentage are gram-positive or spore forming). However, the addition of the gene datasets allows greater insights into the functionality of their microbiome samples. This can improve a researcher or clinicians ability to understand their data and help direct more targeted research questions as they explore their samples and unique microbiomes.

## Statistical Annotations of Taxa Sets

Moreover, we provide an interface for identifying differential features between sets of taxa. Given two lists of taxa, possibly derived from microbiome data, a user may be interested to know whether any of the annotations in TMD are significantly different between the two groups. We provide an interface for this analysis. For annotation features which are numeric, we gather all features for taxa in both lists and perform a t-test. For categorical features we perform a chi-square test. If the input data includes relative abundance with the associated taxa, our interface can also process this information. For this purpose we generate a list of annotation values based on randomly sampling in proportion with the relative abundance of each microorganism then test the sampled lists above.

We used samples from the MetaSUB project (7), to test the statistical annotations of The Microbe Directory v2.0. MetaSUB project aims to catalog the microbiomes of cities around the world. Using a taxonomic set of 3,699 samples from 58 cities, we evaluated whether TMD v2.0 could provide interesting and informative ecological and phenotypic data for each sample in this data set.

We explored two hypotheses for Bacteria and Eukaryote: i) That a cities climate would affect the fraction of species which could form spores, ii) That a cities proximity to the coast would increase the fraction of halotolerant species in the samples. Using our taxonomic annotations and metadata from MetaSUB (i.e. Type of climate), we plotted the values of Spore-Forming species by Climate (Figure 3A) and Halotolerant species by coastal proximity (Figure 3B). To statistically evaluate the distribution, we performed two-way ANOVA for the desired trait blocking on the city annotation. Both relationships were found to be significant. Climate affected the fraction of spore forming species with *p*<2e-16. Likewise, Coastal proximity affected halotolerance with *p*=3.36e-12. In both cases the city (our control factor), also had significant relationships with *p*<2e-16 in both cases.

**Figure 3:**
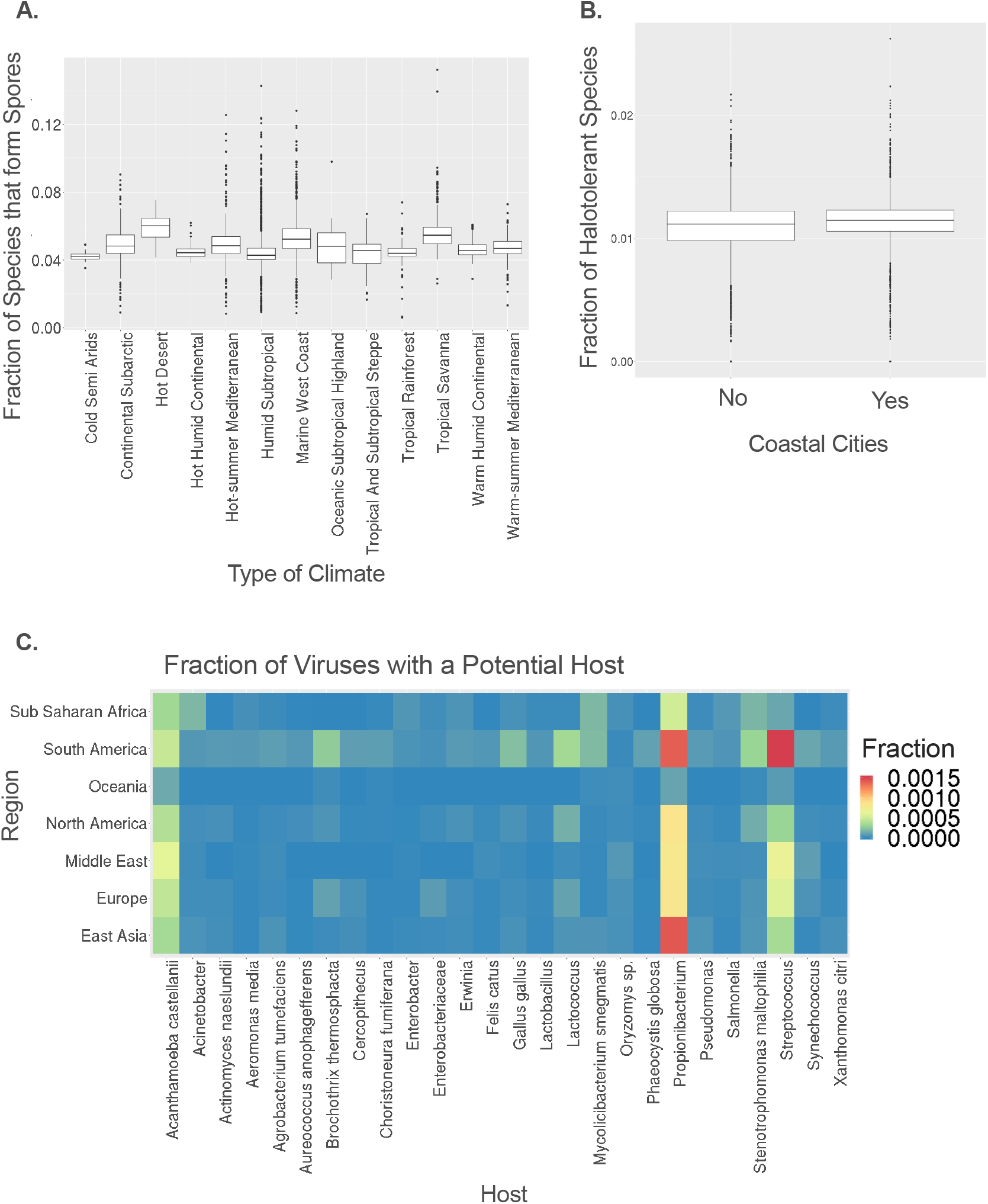
Statistical annotation analysis. Data set from MetaSUB project. **A**.Fraction of species that form spores on the different climate types. **B**.Fraction of Halotolerant species in coastal cities. **C**. Fraction of viruses with a potential host.

We also wanted to explore which were the most common virus-hosts in different regions given the virus annotation from the MetaSUB dataset. We found that the most common hosts are two Bacteria: *Propionibacterium* and *Streptococcus*, and the Eukaryote *Acanthamoeba*. Interestingly, two animals were retrieved as hosts from this analysis: The domestic cat *Felis catus* and the common chicken *Gallus gallus*. These results coincide with the nature of the data, since the MetaSUB study aims to study the urban microbiomes and these taxa are cataloged as human commensals and free living organisms (8; 11).

## Community Based Survey for Correction and Annotation of Taxa

One of the major principles behind The Microbe Directory, was building an open-access resource by researchers for researchers. Thus, one of the goals of expanding TMD to version 2.0, was to build an interface that would allow individuals to contribute with corrections and taxa annotations. Users can contribute through the open survey from KoBoToolbox (Figure 4). This survey is available in English and Spanish to make TMD accessible to more people. This data will be further curated and uploaded to the web interface.

**Figure 4:**
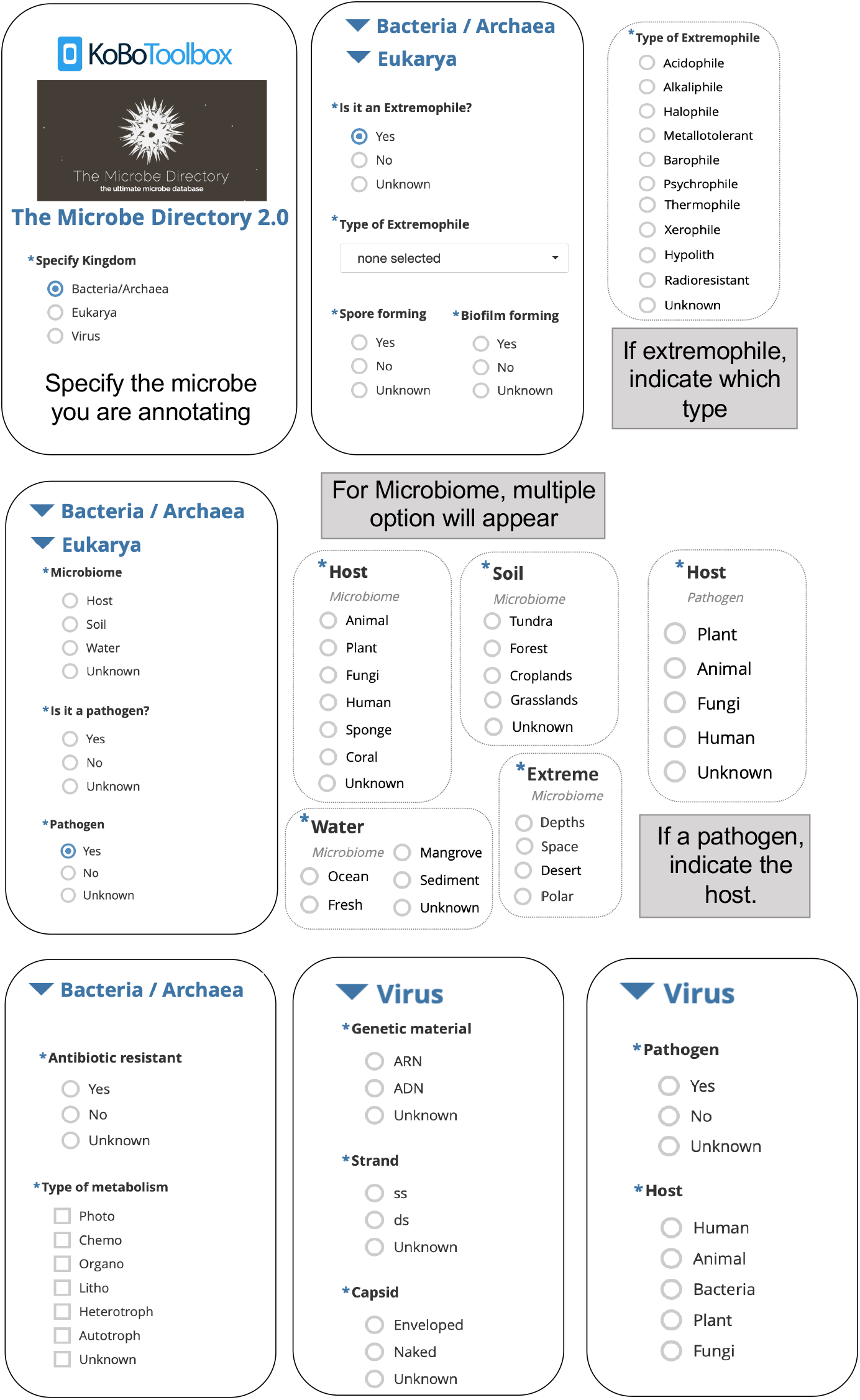
KoboToolbox interface (English version). The survey allows users to contribute with microbial annotations and phenotypic features based on the domain schema of The Microbe Directory v2.0

## Data Availability and Contribution

A web interface for the Microbe Directory may be found at https://microbe.directory. Source code for collating and cleaning tables along with a CSV spreadsheet version of the database may be found at the GitHub repository https://github.com/dcdanko/MD2. The curated gene sets may also be found at this repository. The open survey to contribute new microbes and annotations may be found at the KoBoToolbox. Additional annotations and corrections may also be found submitted at the GitHub.

We note that users of the Microbe Directory should credit the constituent databases relevant to their work. While TMD is intended to provide an accessible and consistent interface to these databases, it in no way precludes the significant amount of work which was required to build these databases.

## Acknowledgements

We would like to thank Naveed Shah and all the students-researchers who helped curate the data for the original Microbe Directory without which this project would never be possible. We would also like to thank the Epigenomics Core Facility at Weill Cornell Medicine, funding from the Irma T. Hirschl and Monique Weill-Caulier Charitable Trusts, Bert L and N Kuggie Vallee Foundation, the WorldQuant Foundation, The Pershing Square Sohn Cancer Research Alliance, NASA (NNX14AH50G, NNX17AB26G), the National Institutes of Health (R25EB020393, R01NS076465, R21AI129851), the Bill and Melinda Gates Foundation (OPP1151054), and the Alfred P. Sloan Foundation (G-2015-13964).

